# Flower visitation through the lens: Exploring the foraging behaviour of *Bombus terrestris* with a computer vision-based application

**DOI:** 10.1101/2024.07.10.602888

**Authors:** Zsófia Varga-Szilay, Gergely Szövényi, Gábor Pozsgai

## Abstract

To understand the processes behind pollinator declines, and thus to maintain pollination efficiency, we also have to understand fundamental drivers influencing pollinator behaviour. In this study, we aim to explore the foraging behaviour of wild bumblebees, recognizing its importance from economic and conservation perspectives. We recorded *Bombus terrestris* on *Lotus creticus*, *Persicaria capitata*, and *Trifolium pratense* patches in five-minute-long slots in urban areas of Terceira (Azores, Portugal). For the automated bumblebee detection, we created computer vision models based on a deep learning algorithm, with custom datasets. We achieved high F1 scores of 0.88 for *Lotus* and *Persicaria*, and 0.95 for *Trifolium*, indicating accurate bumblebee detection. We found that flower cover per cent, but not plant species, influenced the attractiveness of flower patches, with a significant positive effect. There were no differences between plant species in the attractiveness of the flower heads. The handling time was longer on the large-headed *Trifolium* than those on the smaller-headed *Lotus* and *Persicaria*. However, our result did not indicate significant differences in the time bumblebees spent on flowers among the three plant species. Here, we also justify computer vision-based analysis as a reliable tool for studying pollinator behavioural ecology.

## 1. Introduction

In the Anthropocene, the populations of native pollinators are declining worldwide (Nath et al., 2023; Potts et al., 2010), posing a threat to pollinator-dependent crops and wildflowers (Biesmeijer, 2006; Vanbergen & Initiative, 2013). This, in turn, hampers ecosystem functioning of natural and agroecosystems and jeopardises the delivery of ecosystem services vital for humans. While key pressure on pollinators influencing this decline, such as climate change (Kerr et al., 2015; Martinet et al., 2020), land-use change, habitat loss or fragmentation (Vanbergen, 2014), and pesticide inputs (Godfray et al., 2014; Stanley et al., 2015), are well studied (Dicks et al., 2021), their effects on the behaviour of pollinators remain unexplored. Indeed, the effective composition of pollinator communities not only changes with the disappearance of species but also with the subtle behavioural changes, that often go undetected, as the remaining species adapt to new conditions (Kaiser-Bunbury et al., 2010; Schweiger et al., 2010). Since these changes can negatively impact behaviour-mediated ecosystem functions such as pollination (Lippert et al., 2021), to slow down or stop negative population trends and promote recovery we need to understand previously unexplored and intricate details of plant-pollinator interactions, including those related to pollination behaviour (Burkle & Alarcón, 2011; Byers, 2017).

In addition to the undeniable importance of domesticated honeybees (*Apis* sp.) in farming, native and domesticated bumblebees (*Bombus* sp.) play a significant role not only in pollinating wildflowers in natural ecosystems but also in agricultural crop production by maintaining high yields (Rao & Stephen, 2009; Velthuis & Doorn, 2006). Indeed, among wild bees, bumblebees are the highest contributors to crop pollination (Kleijn et al., 2015; Ollerton et al., 2011). Yet, they are particularly exposed to the effects responsible for global pollinator declines (Goulson et al., 2008; Soroye et al., 2020; Williams & Osborne, 2009), and their populations have also been reported to show steep declining trends which were predicted to continue and even accelerate for many species (Ghisbain et al., 2023; Nieto, 2014). This decline is exacerbated by changes in bumblebee behaviour due to various anthropogenic and biotic impacts. For instance widely used agrochemicals and viral or parasite infections can cause non-lethal changes (Varga-Szilay & Tóth, 2022), such as reduced homing ability, colony growth (Stanley et al., 2016), and food intake capacity (Feltham et al., 2014) of the individuals, through which the existence of entire colonies can be jeopardized.

One of the most important behaviours likely to be prone to changes is foraging, which is vital in driving the fitness of bumblebee populations, and thus long-term pollination efficiency. Since it is of high economic and conservation importance, numerous studies scrutinised this behaviour and found that the foraging success and homing ability of bumblebees can be influenced by a multitude of factors. Large-scale abiotic factors such as temperature, humidity, or pesticide exposure could affect flower handling and spatial foraging behaviour (Gill et al., 2012; Samuelson et al., 2016). Small-scale external factors include flower morphology and colour, the characteristics of flowering patches, parasites (Gillespie & Adler, 2013), the cost of flight between patches (Goulson, 2000), forager density, and the spatial distribution of flowers (Geslin et al., 2014). Internal factors, such as learning abilities (Evans et al., 2017) and physical conditions of foragers (Gegear et al., 2006), also play a role. Despite these influences, bumblebees show high behavioural plasticity and forage efficiently under a wide range of environmental conditions (Goulson, 2010; Jha & Kremen, 2013; Zimmerman, 1981) making them less constrained in their foraging behaviour than other insects (Goulson, 2010). To appropriately assess anthropogenic impacts, though, a mechanistic understanding of fundamental foraging behavioural patterns, such as how resource handling times and visiting frequencies depend on plant species or flower density-dependent carrying capacity of patches is needed (Jha & Kremen, 2013). Yet, key information on these important aspects of bumblebee ecology is scarce, most likely because of the laborious means of data collection. In fact, whilst there is a large set of tools for studying pollinator community ecology, high-throughput methods for effectively monitoring behavioural changes of wild bumblebee populations are still missing, which substantially hamper study efforts.

Traditionally used observational methods for recording pollinators’ behaviour and activity, such as transects, mark-recapture, and timed count-based observations, are not only time-consuming and difficult to standardise (Darras et al., 2019) because they highly depend on the skills of the person conducting the field observations, but they also can disturb the insects’ natural behaviour and thus skew the result (Besson et al., 2022; Tuia et al., 2022). Although standardised behavioural observations under laboratory conditions, at least partially, address these issues, they are challenging to adapt to wild conditions, especially because these studies are often conducted on commercially produced bumblebees rather than wild populations (Treanore et al., 2021).

However, modern methods, such as using video recording to observe the foraging behaviour of insect floral visitors around flower sources present an opportunity for increasing efficiency, decreasing disturbance, and saving human labour. When combined with state-of-the-art technologies, including computer vision and deep learning techniques, video recordings can provide novel solutions to species identification (Bjerge et al., 2023; Spiesman et al., 2021) and monitoring communities (Besson et al., 2022) or insect pests (Preti et al., 2021). Indeed, these methods are increasingly used in pollination ecology to detect flower-visiting insects (e.g. (Ratnayake et al., 2022; Sittinger et al., 2024)) and recent trends suggest that the future of pollinator research is going to be shaped by the use of artificial intelligence (AI)-based tools (Barlow & O’Neill, 2020; Høye et al., 2021; van Klink et al., 2022). Despite their numerous advantages, few studies use these methods to monitor pollinators in real-time (but see (Bjerge et al., 2021; Ngo et al., 2021)) and even less to examine unmarked insects behaviour outdoors, under natural or near-natural conditions (but see (Bjerge et al., 2022; Ratnayake et al., 2021)). One of the reasons may be that, whilst the application of AI tools rarely needs any specialised technological hardware, at their current development stage, these methods often require specific computational skills, which extend beyond the expertise of most ecologists. The intensive use of a non-ecology-related discipline thus highlights the need for enhancing collaboration between ecologists and computer scientists (Carey et al., 2019).

Additionally, the effectiveness of computer vision and object detection in field settings, and thus the success of data extraction, depends on several environmental factors such as light conditions, wind strength, and precipitation. To overcome these challenges, some studies used artificial platforms (Sittinger et al., 2024) or fixed flower heads to sturdy surfaces (Steen, 2017). However, no studies have yet been conducted to assess pollinator behavioural patterns using computer vision in completely uncontrolled field settings.

In this research, we used computer-vision-based methods to study wild populations of buff-tailed bumblebees (*Bombus terrestris* (Linnaeus, 1758), Hymenoptera, Apidae) foraging on three wild-growing, insect-pollinated plants: Cretan bird’s-foot trefoil (*Lotus creticus* Linnaeus, 1753, Fabales, Fabaceae), pink-headed knotweed (*Persicaria capitata* (Buch.-Ham. ex D.Don) H.Gross, Caryophyllales, Polygonaceae), and red clover (*Trifolium pratense* L., Fabales, Fabaceae) in urban areas. This work aimed to explore and understand the foraging behaviour of bumblebees under field conditions and to describe the differences between their behavioural patterns on the different flower species and with varying flower densities. At the same time, our alternative objective was to test whether video-based recording methods combined with computer vision-based analysis are suitable for exploring bumblebee behaviour under field conditions.

We aimed to answer the following questions:

- Question 1: How many bumblebees a flower patch can support within a specified time unit? Is there a difference in the visitation of flower patches per time unit?
- Question 2: How many bumblebees can occupy a flower patch simultaneously?
- Question 3: What proportion of their total time (‘*bumblebee-time’*) bumblebees spend on flowers (handling time) compared to time spent on non-flowery areas (travelling time)?

Our hypothesis was that the assumed optimal foraging behaviour of the nectar- and pollen-gathering bumblebees show significant differences in the time spent on flowers among the three plant species and that it is adjusted to the characteristics of the resources (three types and density of flower patches and flower head size of plants).

## 2. Material and methods

### 1.1. Study sites

The study was conducted on Terceira Island (Azores, Portugal) between May and September 2022. The sampling sites were located in urban areas at 38°48’06.0” N, 27°15’17.8” W, 38°44’14.7” N, 27°16’07.8” W, and 38°47’38.7” N, 27°15’24.0” W (**Figure S1**). We recorded the bumblebees (*Bombus terrestris*) on Cretan bird’s-foot trefoil (*Lotus creticus*; indeterminate biogeographic origin), pink-headed knotweed (*Persicaria capitata*; introduced invasive), and red clover (*Trifolium pratense*; introduced naturalised) patches.

### 1.2. Data collection

Videos were recorded with GoPro Hero9 action cameras in approximately five-minute-long slots in 5K resolution (5120 by 2880) at 30 frames per second speed on 60×60 cm square quadrats (Figure 1). We excluded videos without bumblebees from the analysis (nine videos from *Lotus* and two from *Persicaria*), which resulted in 15, 18, and 15 videos from *Lotus, Trifolium,* and *Persicaria*, respectively.

**Figure 1:**
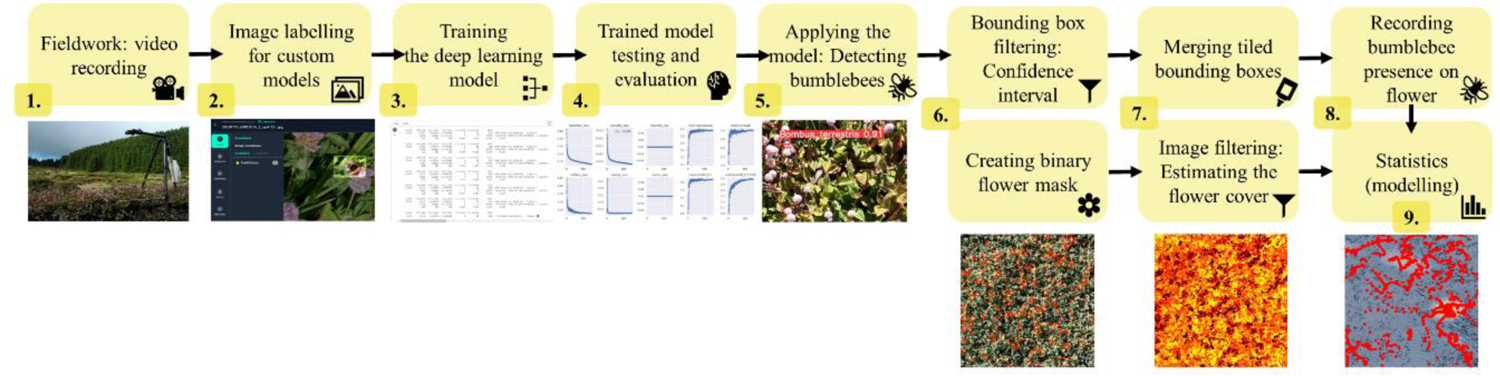
Flowchart of the data collection, preparation and analysis to assess bumblebee behavioural differences of three plant species.

Although we attempted to take the videos at the same height from the ground, this was not always possible due to the uneven surface. Therefore, to allow size and area estimations, each setup was calibrated with a millimetre-precision scale and the real-life length of one pixel was calculated. Metadata showing recording location, date and time was linked to each video and used in the analysis. Temperature (C°) and humidity (%) were measured, and wind strength (Beaufort Wind Scale) and cloudiness (direct sunshine, overcasted, or cloudy) were estimated on-site.

### 1.3. Data processing

Videos were split into frames and frame-level information was further used for training and analysis. For labelling the training set for the deep learning algorithm, three-second segments (90 frames) were cut from the beginning of each video or from the frame where the first bumblebee(s) appeared in the video. The images were manually annotated by drawing bounding boxes of one label class (‘bumblebees’) around the bumblebees with the help of Roboflow annotation tool (Dwyer et al., 2024). The datasets (*Lotus*: 4308 images (Varga-Szilay, 2023b), *Persicaria*: 1908 images (Varga-Szilay, 2023a), *Trifolium*: 2099 images (Varga-Szilay, 2023c)) comprising annotated bumblebees, were split into training, cross-validation, and test sets (as 70, 20, and 10% proportions, respectively, **Table 1**). A proportion of 5% of the original image number was also added as false positive images for each type of image set (**Table 1**).

**Table 1:** The dataset of images used to train the object detection model (YOLOv5). The table shows the number of ‘bumblebee’-labelled images, as well as the false positive images (e.g. with background or shadows) without label.

| Type of set | Type of images | Flower species |  |  |
| --- | --- | --- | --- | --- |
|  |  | *Lotus* | *Trifolium* | *Persicaria* |
| Test | Labeled | 387 | 254 | 202 |
|  | False positive | 19 | 12 | 10 |
| Train | Labeled | 2889 | 68 | 1247 |
|  | False positive | 144 | 1354 | 62 |
| Validation | Labeled | 828 | 391 | 369 |
|  | False positive | 41 | 20 | 18 |
| **Total** |  | **4308** | **2099** | **1908** |

For training, to keep the resolution high yet allow tiling with 640-pixel (px) segments, the images were expanded from 5120×2880 to 5120×3200 and then they were cropped onto 640 x 640 px tiles (pre-processing). For the automated detection of bumblebees in the videos we created deep learning-based computer vision models for each plant species separately, using YOLOv5 (You Only Look Once (Jocher, 2020)) with custom datasets for each plant species. For the training process we used either Google Colab (Tesla 4T 15102MiB GPU) or a desktop PC (11^th^ Gen Intel(R) Core(TM) i9/11900KF @ 3.50GHz, 64 GB RAM, 8 Cores, Win11, NVIDIA GeForce RTX 3060, 12288 MB) with a PyTorch (Paszke et al., 2019) environment with 0.01 learning rate (LR). All models were trained to 300 epochs with 64 batch size and default hyperparameters. See the specific evaluation metrics of YOLOv5 models in **Figure S2.** Further analysis was performed with a data set (bounding box set) filtered twice to minimize false bumblebee detections. For bounding box filtering, a confidence level of 0.7 was used for the object detection results for *Trifolium*, and 0.8 for *Lotus* and *Persicaria*. To avoid multiple detections of one bumblebee through the tiling process after the tiles were merged we calculated the Euclidean distance between each bounding box and merged the boxes if the distance between them was less than or equal to the size of half of an average bumblebee (15 mm calculated from the calibrated pixel sizes) (post-processing). To test the post-processing accuracy of the model on filtered bounding boxes, we randomly selected five frames from five videos for each flower species and compared the manual detections with those predicted by the model (**Table S1**).

To crop the quadrat from the video, the upper left corner of the physically placed square quadrat was digitally identified, and the *x1* and *y1* coordinates of this corner were recorded. Then, the remaining coordinates (*x2, y2*) of the 60 x 60 cm square were computed knowing the one pixel/mm value. The quadrats were then extracted from each video frame based on these calculated coordinates. Bumblebee detections were only kept if the centroids of the bounding boxes were within the quadrat.

For flower detection, we manually determined the flower species-specific upper and lower Hue, Saturation and Brightness (HSB) colour thresholds from the unedited images (Figure 2a) with the ImageJ software (Rasband, 2018)) (*Lotus*: 11, 42, 120, 255, 160, 255; *Persicaria*: 140, 255, 0, 140, 40, 255; *Trifolium*: 85, 255, 0, 247, 186, 255) and used these in a colour filtering process. The ImageJ-compatible HSB ranges were later converted to Python-compatible Hue, Saturation and Brightness (HVS) colour thresholds. To calculate an average flower colour within this range, the ‘optimal flower colour’ (OFC), ten random frames were selected from each video and a colour mask for the HVS colour thresholds was applied to each of these frames (Figure 2b). Then the median of all of these colour masks was determined for each flower species. To estimate the per cent cover of flowers (henceforth flower cover) in the quadrat, we created a binary mask based on the HSV range, counted the number of masked pixels and then the proportion of flower area in a quadrat was calculated by dividing this number by the total number of pixels in the quadrat. The value was calculated from one random frame for each video separately.

**Figure 2:**
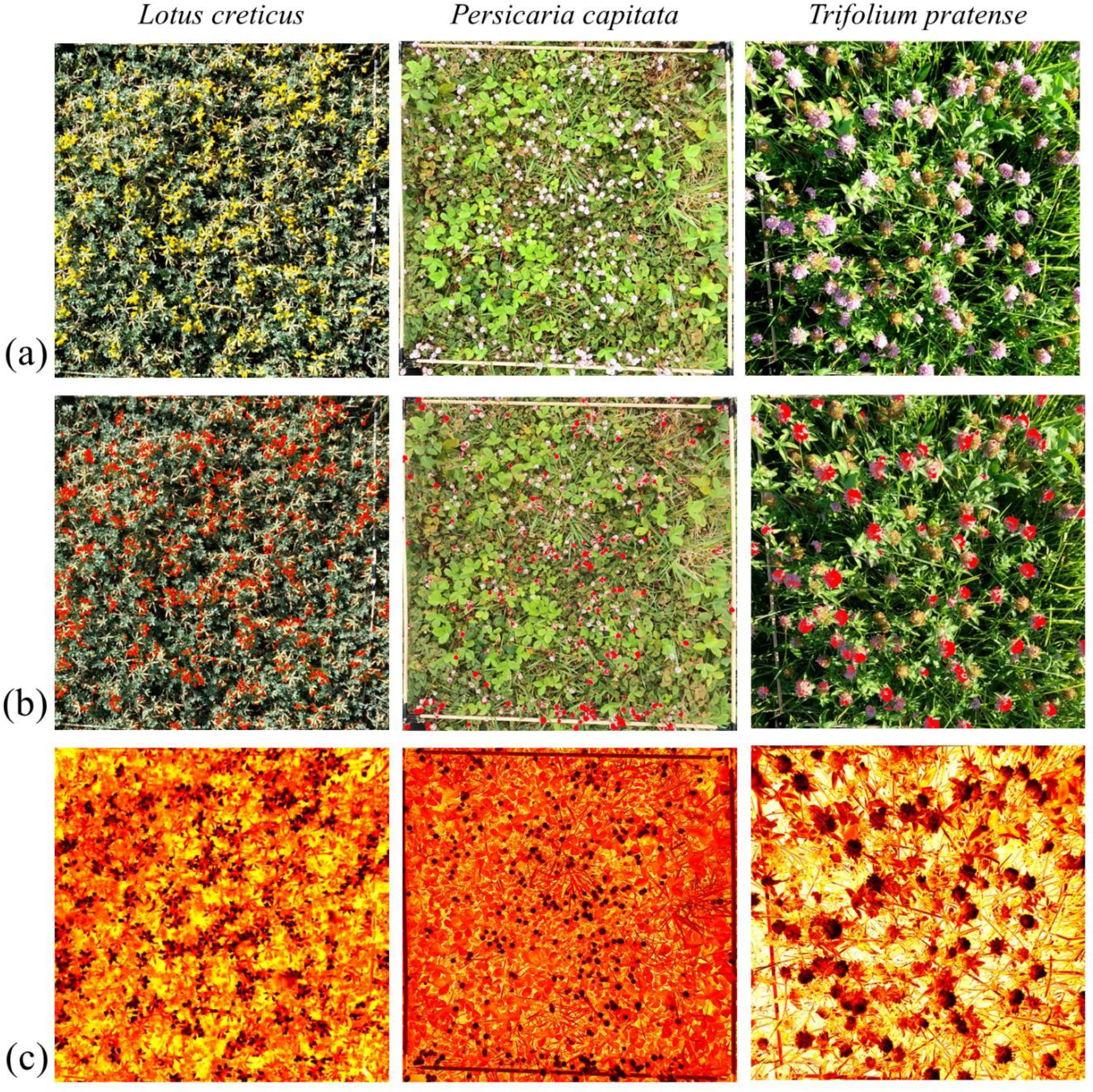
Example of (a) original images, (b) images filtering with binary mask, and (c) heatmap-based images indicating the Euclidean distance of the actual colour from the ‘optimal flower colour’ (brighter red indicates greater distance).

To create the heatmap, we utilized average similarity from the OFC of 3000 frames per video. To get this colour similarity from the predefined OFC we calculated the distance between each pixel in an HSV image (= one frame) and the OFC by extracting the Hue, Saturation, and Value components, then computing the Euclidean distance, taking into account the circular nature of the Hue component (Figure 2c). The areas of bounding boxes marking bumblebees were excluded from the average calculation.

### 1.4. Statistical analysis

To visualise the frequency of pixel colour distances from the optimal flower colour on a histogram for the flower species, we took a random sample (100,000 points or as many as were available) from areas on the heatmap where bounding boxes were and from the whole heatmap (full quadrat).

To determine how many bumblebees were attracted to each flower patch (for **Question 1**), we calculated the sum number of bumblebees per video divided by the total number of frames in the video and used this measure as a proxy for the attractiveness of a flower patch. When this number was standardized by the flower cover too (assuming that bees are more likely to handle flowers when flower sources are more abundant) we got a proxy for the carrying capacity of each plant species (for **Question 2**).

To determine whether bumblebees were on flowers, we used the flower colour thresholds to mask the bounding box area enclosing a bumblebee from a frame in which the animal was not present (to see the flower colours rather than those of the bumblebee’s) and calculated the proportion of these masked pixels relative to the total pixels of the bounding box area. If the masked pixels (indicating flower colours) reached 20%, we considered the bumblebee to be on a flower at the moment of the detection. Thus, we refer to ‘handling time’ or ‘on-flower time’ when the detected bumblebee was on a flower, while bumblebee detections, where the insect was unlikely on a flower, are termed ‘travelling time’ or ‘off-flower time’. To estimate a proxy for handling time (for **Question 3**), we calculated the proportion of ‘bumblebee-on-flower-time’ (summarising all time units (frames) that all detected individuals cumulatively spent on flowers) to all ‘bumblebee-time’ (that summarised all time units that all individuals, cumulatively, spent within the quadrat).

We used simple linear regression analysis and linear mixed models (wind strength and temperature as random effects) to evaluate the impact of plant species and flower cover on bumblebee behaviour. Mixed models were kept when they demonstrated superior performance compared to the simple linear models, in terms of the Akaike Information Criterion (AIC), and the fit was not singular, otherwise, the simple linear regression models were used. All models were tested for homoscedasticity and the normality of residuals using the Shapiro-Wilk and the Goldfeld-Quandt tests, respectively. When assumptions of normality and homoscedasticity were not met, we either square-root or cubic-root transformed the response variables (**Table 2**). Heteroscedasticity was accounted for by applying heteroscedasticity-robust standard errors with the help of the ‘sandwich’ R package.

**Table 2.:**
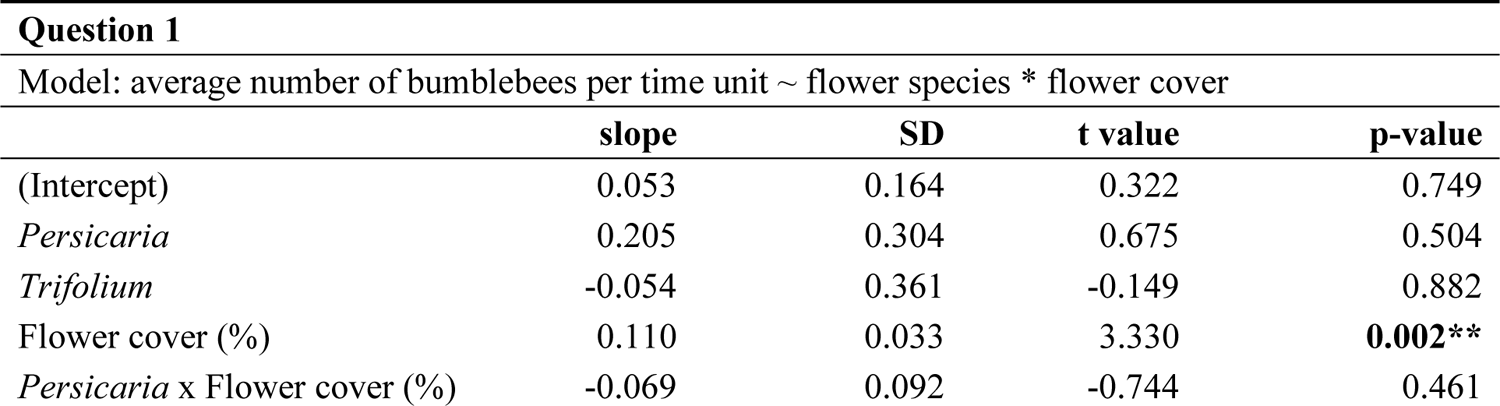
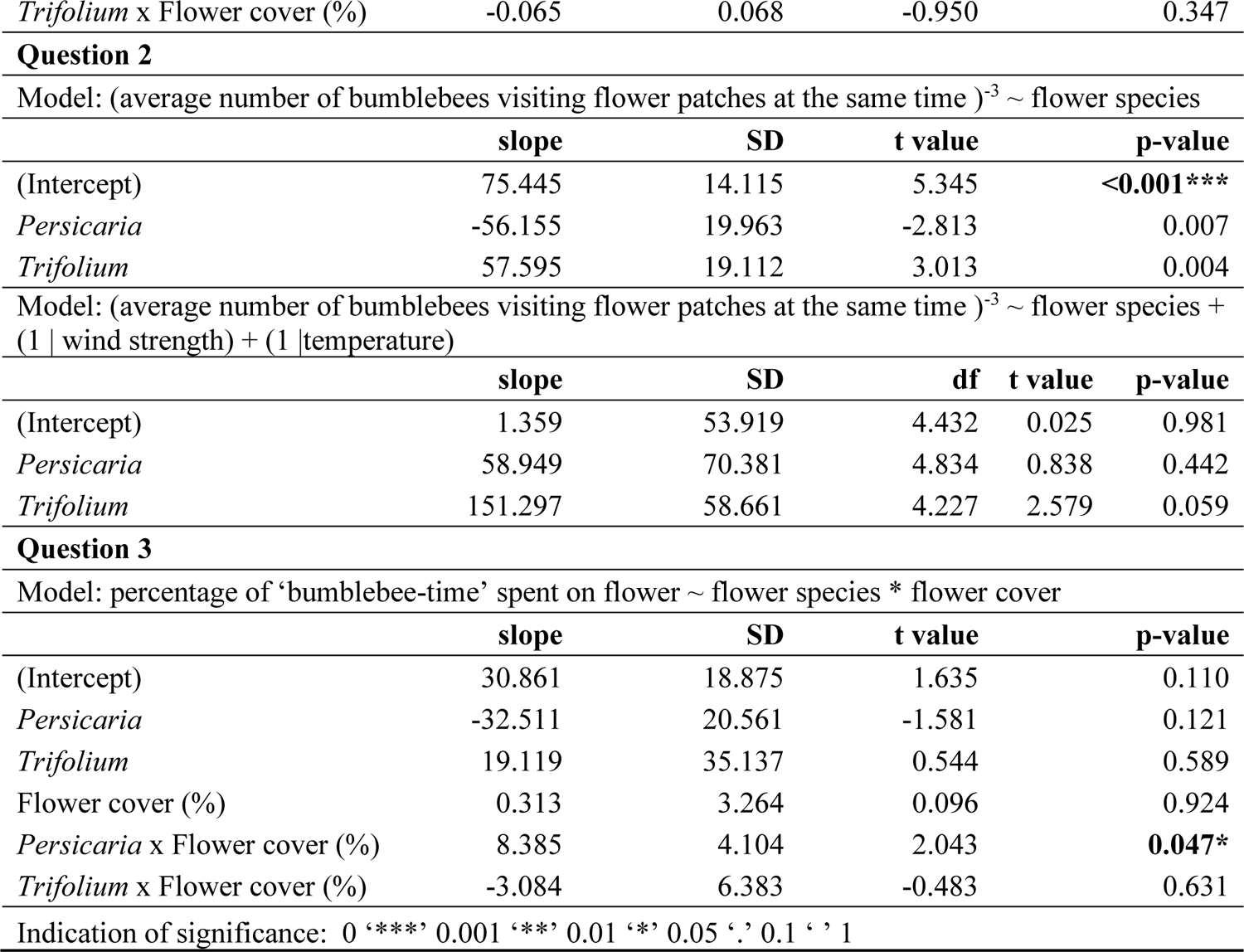
Summary statistics of the linear regression and linear mixed models.

| Question 1 |  |  |  |  |
| --- | --- | --- | --- | --- |
| Model: average number of bumblebees per time unit ~ flower species * flower cover |  |  |  |  |
|  | slope | SD | t value | p-value |
| (Intercept) | 0.053 | 0.164 | 0.322 | 0.749 |
| *Persicaria* | 0.205 | 0.304 | 0.675 | 0.504 |
| *Trifolium* | -0.054 | 0.361 | -0.149 | 0.882 |
| Flower cover (%) | 0.110 | 0.033 | 3.330 | **0.002\*\*** |
| *Persicaria* x Flower cover (%) | -0.069 | 0.092 | -0.744 | 0.461 |
| *Trifolium* x Flower cover (%) | -0.065 | 0.068 | -0.950 | 0.347 |

Question 2 Model: (average number of bumblebees visiting flower patches at the same time)<sup>-3</sup> ~ flower species
|  | slope | SD | t value | p-value |
| --- | --- | --- | --- | --- |
| (Intercept) | 75.445 | 14.115 | 5.345 | <0.001*** |
| *Persicaria* | -56.155 | 19.963 | -2.813 | 0.007 |
| *Trifolium* | 57.595 | 19.112 | 3.013 | 0.004 |

|  | slope | SD | df | t value | p-value |
| --- | --- | --- | --- | --- | --- |
| (Intercept) | 1.359 | 53.919 | 4.432 | 0.025 | 0.981 |
| *Persicaria* | 58.949 | 70.381 | 4.834 | 0.838 | 0.442 |
| *Trifolium* | 151.297 | 58.661 | 4.227 | 2.579 | 0.059 |

Question 3 Model: percentage of 'bumblebee-time' spent on flower ~ flower species \* flower cover
|  | slope | SD | t value | p-value |
| --- | --- | --- | --- | --- |
| (Intercept) | 30.861 | 18.875 | 1.635 | 0.110 |
| *Persicaria* | -32.511 | 20.561 | -1.581 | 0.121 |
| *Trifolium* | 19.119 | 35.137 | 0.544 | 0.589 |
| Flower cover (%) | 0.313 | 3.264 | 0.096 | 0.924 |
| *Persicaria* x Flower cover (%) | 8.385 | 4.104 | 2.043 | 0.047* |
| *Trifolium* x Flower cover (%) | -3.084 | 6.383 | -0.483 | 0.631 |

Indication of significance: 0 '\*\*\*' 0.001 '\*\*' 0.01 '\*' 0.05 '.' 0.1 ' ' 1

For data preparation and visualization, we used Python-3.8.16 (Python Software Foundation, 2019) environment and torch-1.13.1, with the help of ‘NumPy’ version 1.23.5 (C. R. Harris et al., 2020), ‘Pandas’ version 1.5.3 (The pandas development team, 2022), ‘cv2’ version 4.7.0 (Bradski, 2000), ‘ffmpegcv’ (FFmpeg Developers, 2016), ‘matplotlib’ version 3.7.1 (Hunter, 2007), and ‘scipy’ version 1.10.0 (Virtanen et al., 2020) libraries. For data preparing, modelling and the visualisation of model results, we used the ‘dplyr’ (Wickham et al., 2023), ‘readr’ (Wickham et al., 2024), ‘purrr’ (Wickham & Henry, 2023), ‘lmtest’ (Zeileis & Hothorn, 2002), ‘sandwich’ (Zeileis et al., 2020), and ‘ggplot2’ (Wickham, 2016) packages in an R environment (R version 4.4.0 (R Core Team, 2021)).

## 3. Results

We recorded 134,865 frames in 15 videos on *Lotus*, 154,151 frames in 18 videos on *Trifolium*, and 129,145 frames in 15 videos on *Persicaria*. The average percentage of flower cover was the lowest on *Persicaria* (2.86%, SD ± 0.85), while it was 4.48% (SD ± 2.23) and 5.29% (SD ± 1.12) on *Lotus* and *Trifolium,* respectively.

Higher F1 scores for *Lotus* and *Trifolium* indicated reliable overall performance of the model in accurately identifying and distinguishing between bumblebees compared to the detection on *Persicaria* (**Figure S3**). After the post-processing, the reliability of the F1 score further increased to 0.88 for the *Lotus* and *Persicaria* and 0.95 for the *Trifolium* (**Table S1**). Overall, the model demonstrated high accuracy in finding the bumblebees and a low rate of false detections. We recorded bumblebees within the quadrats on 177,271 occasions across 418,161 frames. The maximum number of bumblebee individuals at the same time on *Lotus* was 6 (mean = 1.24, SD ± 0.26), on *Persicaria* it was 5 (mean = 1.16, SD ± 0.15), and on *Trifolium* it was 4 (mean = 1.07, SD ± 0.08). There was a difference in the handling time between different plants, with the longest recorded being on *Trifolium* (Figure 3). In addition, bumblebees spent more time off-flower on *Lotus* and *Persicaria* patches than on *Trifolium*.

**Figure 3:**
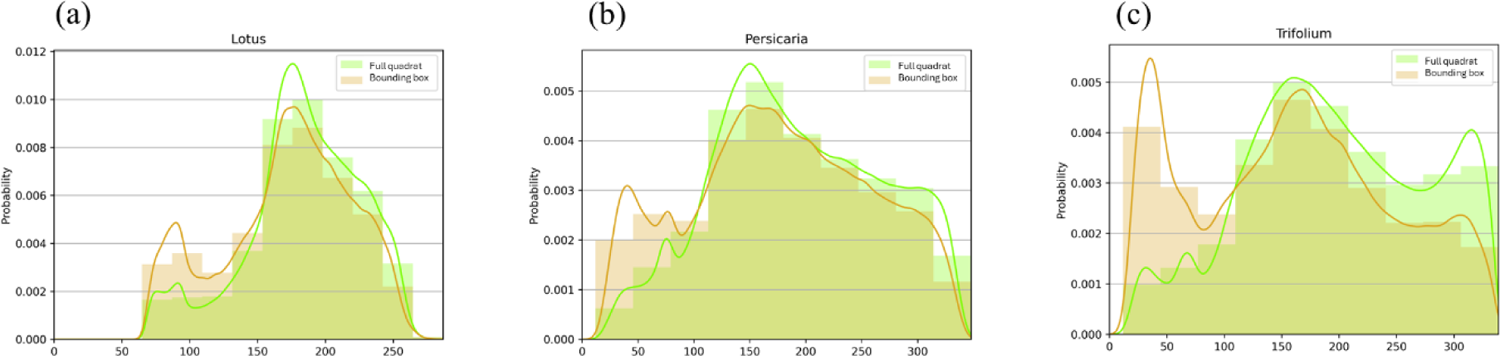
The histograms show the distribution of Euclidean distances of the colours of sample pixels from the predefined optimal flower colour (OFC = 0.0 on the x-axis) on *Lotus* (a), *Persicaria* (b), and *Trifolium* (c) patches. Green coloured bars indicate pixels randomly selected from the full quadrat, whilst the distribution of the colour deviation from OFC under bounding boxes enclosing detected bumblebees is indicated in orange. The colour coding of the smoothed density curves follows that of the bars.

There were more bumblebees in *Lotus* than in the other two plant species within an average time unit. The attractiveness of flowering patches was significantly influenced by flower cover (p < 0.01, **Table 2**) but the plant species did not show significant effects (Figure 4). The curve was steep in the case of *Lotus*, which may have been caused by the few low values of our measure at extremely low cover values. However, the steep relationship between the cover and bumblebee visitation on *Lotus* remained even after those outliers were removed.

**Figure 4:**
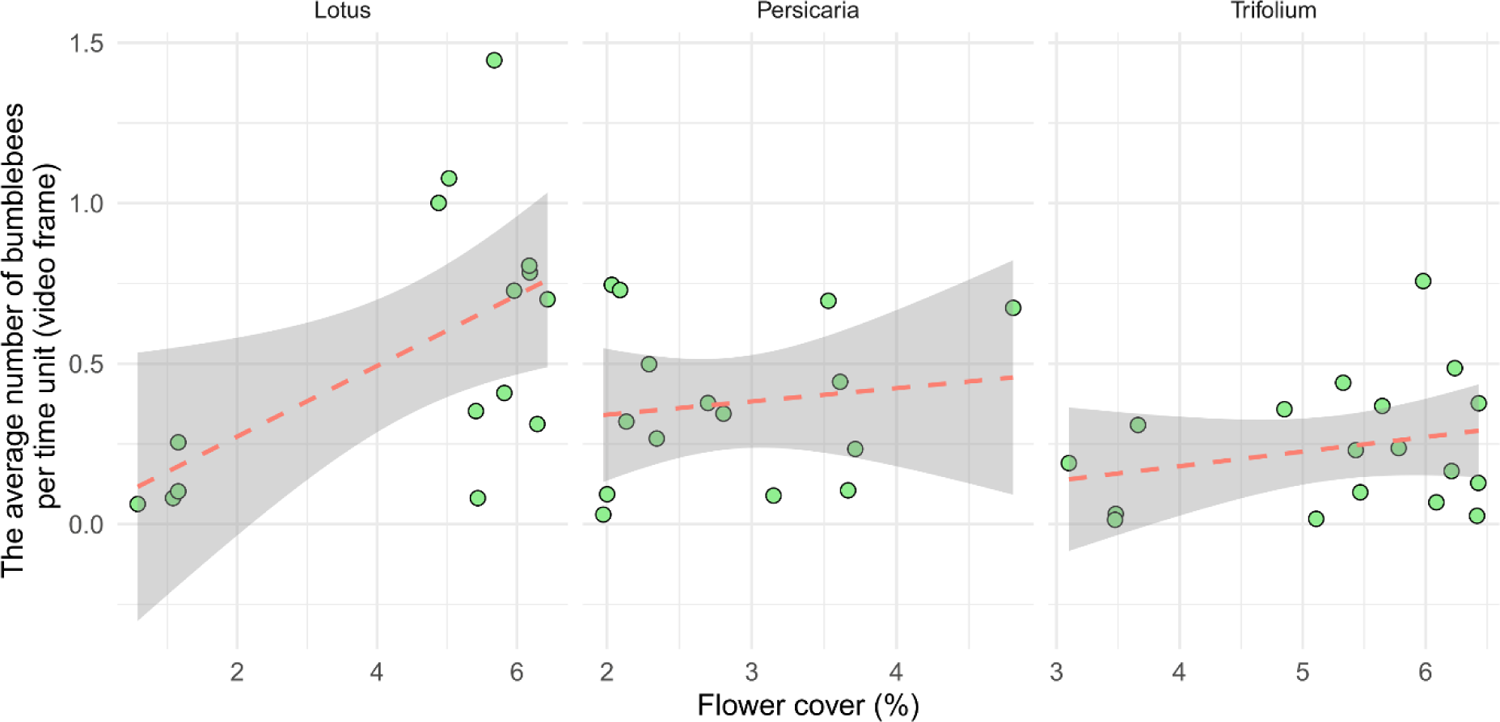
Visitation of flower patches per unit time (video frame) as a function of flower cover separated by plant species. Each point represents one video (n = 15, 15 and 18, for *Lotus*, *Persicaria, and Trifolium*, respectively).

When the plants’ carrying capacity was investigated, the simple linear regression model showed a significant difference (p < 0.001) between the flower species in the average number of bumblebees visiting flower patches at the same time (standardised for flower cover) (Figure 5). However, when we controlled for wind strength and temperature in a random model no significant differences were found between the plant species (**Table 2, Figure S4**).

**Figure 5:**
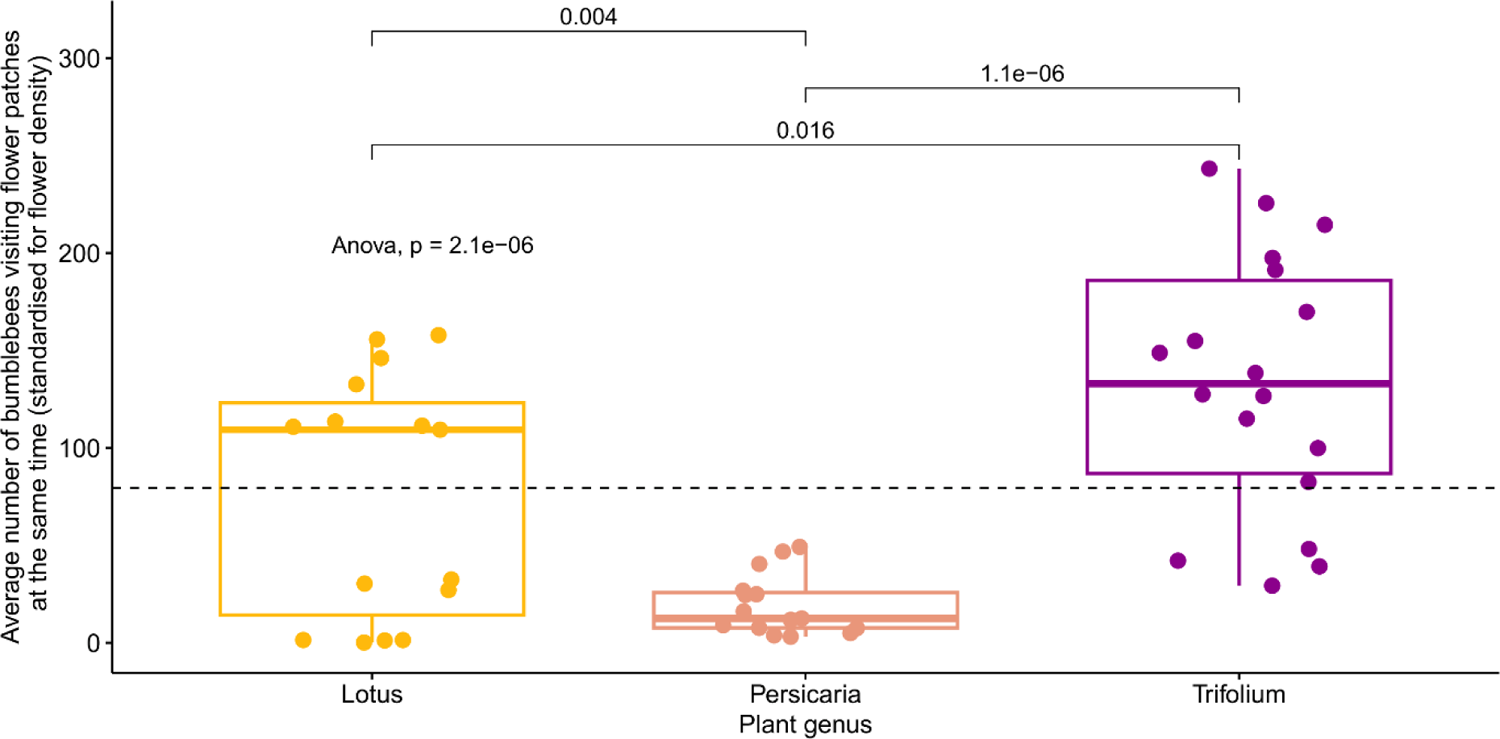
Average number of bumblebees visiting flower patches at the same time (standardised for flower cover) separated by plant species. The global p-value for the ANOVA test is shown in the figure, as are the pairwise comparisons (t-tests) of the averages between plant species. The dashed line shows the mean of the y-axis. Each point represents one video (n = 15, 15, and 18, for *Lotus*, *Persicaria*, and *Trifolium*,respectively).

The model indicated a significant positive correlation between ‘*bumblebee-time*’ (individuals x time spent on flowers) spent on flowers and flower cover but only on *Persicaria* patches (p = 0.047, **Table 2**). There was an indication of a slight but not significant negative interaction between ‘*bumblebee-time*’ spent on *Trifolium* flowers and flower cover (Figure 6).

**Figure 6:**
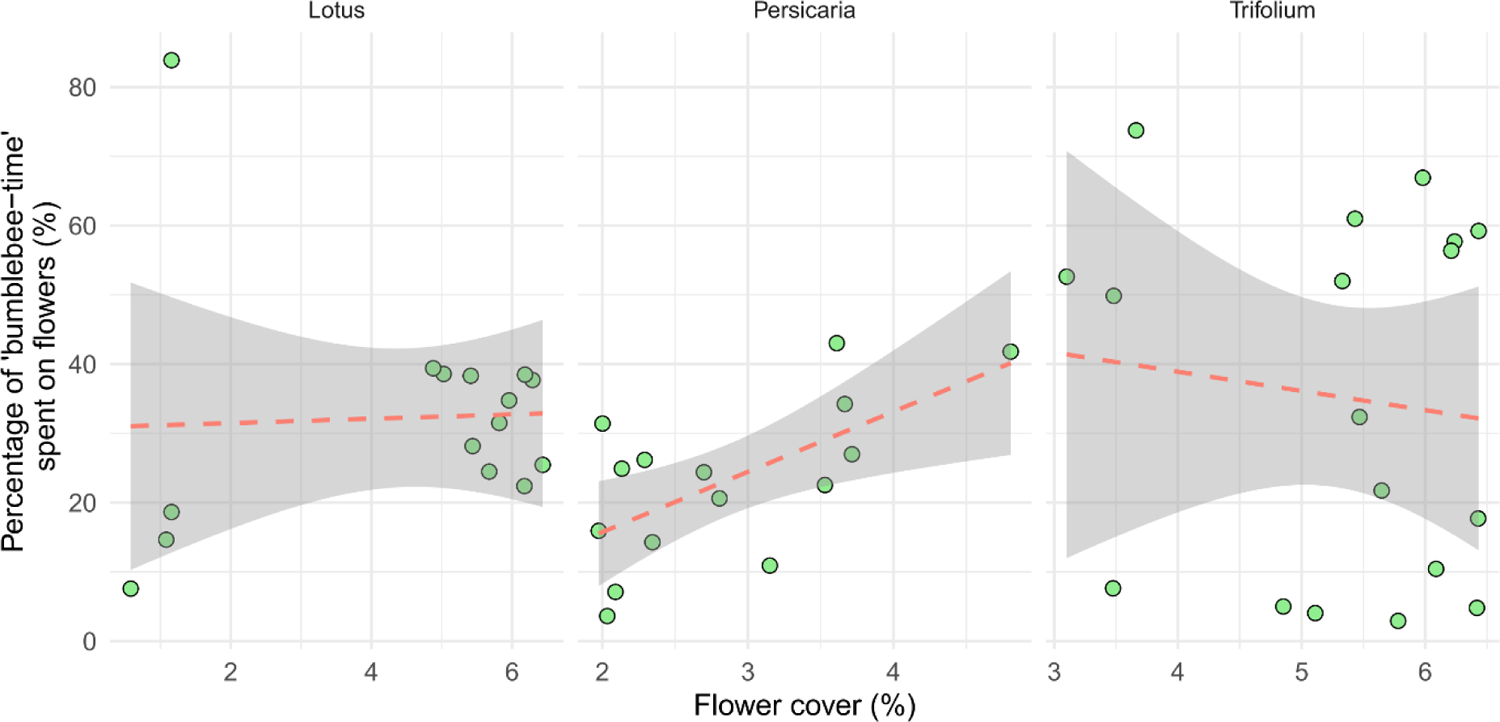
The time bumblebees spent on flowers as a per cent of all time spent in the quadrat (‘bumblebee-time’). Each point represents one video (n = 15, 15, and 18, for *Lotus*, *Persicaria*, and *Trifolium*, respectively).

## 4. Discussion

In this study, we presented a computer-vision-based method to identify differences in the foraging behaviour of wild populations of bumblebees on three insect-pollinated plants among flower species and varying flower cover. We found only slight differences in the foraging behaviour of bumblebees among plant species but detected an indication of the importance of flower cover. This was likely because bumblebees adapted to the characteristics of the flower resources, including flower head sizes, as well as the different flower cover, which influenced their handling time and travelling behaviour.

### 4.1. The attractiveness of the flower patch

We found that the number of bumblebees visiting a patch depended solely on the flower cover and not on the plant species. This aligns with the findings of Vaca-Uribe et al. (2021) (Vaca-Uribe et al., 2021) who reported a positive relationship between the blooming cover and the abundance of insect visitors. In our study, the observed dependency on flower cover was particularly evident with *Lotus*, where patches with low flower cover had fewer bumblebees, while higher flower cover attracted a higher number of individuals, and this relationship between the flower cover and the number of bumblebees remained even after excluding outliers with extreme low cover values. One of the reasons for this is likely that in denser flower patches, bumblebees can spend more time handling flowers, minimizing the energy costs that would be spent searching for a new patch and maximizing resource uptake. However, the exact relationship between increasing cover and bumblebee visitation and its environmental drivers is challenging to determine, as we did not sample flower patches with intermediate flower cover.

Yet, other bumblebee species showed differences in their foraging behaviour, including their patch choice, based on flower cover or flower complexity (Stout et al., 1998).

### 4.2. The carrying capacity of the flower species

The different results from the simple linear and the linear mixed model indicate no clear evidence of whether there is a difference in how many bumblebees can simultaneously occupy each flower species. Indeed, the variability of environmental parameters, such as temperature and humidity, both of which could affect the nectar and pollen production of the plants (C. Harris & Ratnieks, 2024), and the limited sample size may have masked differences among flower species. In addition, other factors such as the risk of predation, the rate of food intake (Pyke, 1984) or the densities of previous and simultaneous foragers (Lázaro et al., 2011) could also be influential, and therefore they should be examined to gain a comprehensive understanding of simultaneous bumblebees patch occupancy. Indeed, based on our field observations, whereas the other two plant species were almost exclusively visited by bumblebees during the recording periods, not only *B. terrestris* visited the *Persicaria* patches but other large-bodies insect taxa, such as flies and other bees, most likely changing bumblebees’ attraction to flowers. Furthermore, the attractiveness of flower heads for foraging bumblebees changes rapidly within a patch, for instance with the amount of available nectar and pollen (Somme et al., 2015) or the speed of rewards replenishment. Additionally, in the case of *Trifolium*, the cultivar can also influence the amount of nectar produced, thereby affecting its attractiveness to visiting insects (Szabo & Najda, 1985). This many sources of variability may explain why we did not detect significant differences between flower species when random variables were included and why results were different when simple regression was used. Thus, although it would be important to determine the influence of plant species on bumblebee carrying capacity, the high dynamism of the system makes this task extremely difficult if not impossible.

### 4.3. The time spent with handling

Bumblebees spend a longer proportion of their total time on flowers (‘bumblebee-time’) relative to non-flower areas on *Trifolium*, compared to *Lotus* and *Persicaria*, likely due to differences in flower head size, resulting in longer handling time on plants with larger heads than on those with smaller ones. However, other factors like flower-specific parameters (e.g. structural complexity (Harder, 1983), nectar concentration and flower depth (Harder, 1986), and the nectar secretion rates (Stout & Goulson, 2002) can also influence how bumblebees optimise behaviour and thus handling time.

The probability of detecting a bumblebee on flower heads was higher on *Trifolium* compared to *Lotus* and *Persicaria*. This was likely because *Trifolium* flower heads are large, requiring a longer handling time, and thus, an increased probability of detecting a bumblebee on flowers. Moreover, the differences in time spent on flowers can also be the result of different strategies bumblebees choose to move between flowers. Since *Lotus* covers the surface in almost two dimensions, bumblebees tend to crawl between flowers (walk from one flower head to the next) but they tend to fly between *Trifolium* flowers, where flower heads are more scattered and vary in flower heights. Indeed, crawling between flowers is preferred as it expends significantly less energy compared to flight (Krell, 2018). Thus, since flying over non-flowery *Trifolium* areas is faster than crawling between flowers in the patches of the other two plants, the shorter off-flower time of bumblebees on *Trifolium* can, at least partially, be explained. Our field observations, especially in the case of *Lotus* with a higher flower cover, supported this theory. Yet, bumblebees may need to make multiple flights between flower heads on the *Trifolium* patch (which is faster), whereas crawling between flower heads results in slower but uninterrupted means of transport on *Lotus*. This suggests that whilst the modes of movement differ, the overall distance covered by foraging individuals between flower heads can be similar. In summary, based on our results, we cannot conclusively support our hypothesis that there are significant differences in the time bumblebees spent on flowers (handling time) among the three plant species. Further research is needed to fully explore the foraging practices of bumblebees and understand how they optimize the time spent handling flowers.

### 4.4. Methodological perspectives

In this study, we also tested the efficiency of a video-based recording method combined with computer vision analysis for studying bumblebee behaviour. Although our results are quite promising and we were able to detect bumblebees under field conditions on all three plant species with high accuracy, the field setting came with several challenges. Shadows, moving backgrounds, and varying light conditions can affect efficient object detection, and lead to false positive detections (Ratnayake et al., 2021). These, however, can be mitigated by improving training datasets, applying post-processing bounding box filters, and background subtraction techniques (Bjerge et al., 2024). Moreover, in natural vegetation, the 3D structure of plants (e.g. varying heights) can cause substantial difficulties.

Furthermore, detecting small insects in high-resolution images remains a challenge for current object detection models, but current advancements (e.g. the latest iteration in the YOLO series, Yolov8 (Jocher et al., 2023) with Slicing Aided Hyper Inference (SAHI) (Akyon et al., 2022)) are enhancing the efficacy of these methods and making image pre-processing (such as tiling) unnecessary. Indeed, in our case, adjusting the camera height to ensure that the quadrat fits within the recorded area while keeping insects recognizable to the model was essential.

Additionally, physical quadrats disturbed foraging bumblebees and interfered with the image analysis (particularly during colour filtering), therefore, at later stages we used digitally designated quadrats, calibrated in post-processing, to avoid these issues. Despite these difficulties, we demonstrated promising results and we are confident that by individually tracking animals, these computer vision-based methods can be effective tools in providing new insight into previously unexplored or controversial behavioural issues of bumblebees.

### 4.5. Future perspectives

The foundational principles of studying bumblebees and other pollinators were established as early as the middle and late 1990s, for example with a boom in research on ‘optimal foraging theory’ (Pyke, 1984). With the recent advancements of AI and computer vision, we now have the opportunity to revisit and improve these foundational studies with greater precision and larger sample sizes without exponentially increasing human labour. These novel technologies can facilitate more comprehensive and accurate studies of behavioural ecology, supporting biodiversity conservation and addressing many of the unanswered questions from earlier research. Nevertheless, further improvements are necessary to enhance the accuracy of insect detection and reliably capture fine-scale changes in behaviour. Studies like ours provide foundational insights that can inform future research without concerns about animal welfare (Lövei & Ferrante, 2024) or promoting cost-effective agricultural practices aligned with sustainability and conservation agriculture (e.g. in selecting plant species for landscape design).

## Author Contributions

Conceptualization, Z.V.S., G.S. and G.P.; Methodology, Z.V.S. and G.P.; Software, Z.V.S. and G.P.; Validation, Z.V.S. and G.P.; Formal Analysis, Z.V.S. and G.P.; Investigation, Z.V.S. and G.P.; Resources, Z.V.S. and G.P.; Data Curation, Z.V.S. and G.P.; Writing – Original Draft Preparation, Z.V.S. and G.P.; Writing – Review & Editing, Z.V.S., G.S. and G.P.; Visualization, Z.V.S. and G.P.; Supervision, G.P. and G.S.; Project Administration, Z.V.S.; Funding Acquisition, G.P., G.S. All authors have read and agreed to the published version of the manuscript.

## Funding

GP was supported by the project Open Access FCT-UIDP/00329/2020-2024 (Thematic Line 1 – integrated ecological assessment of environmental change on biodiversity, https://doi.org/10.54499/UIDB/00329/2020) and by the Azores DRCT Pluriannual Funding (M1.1.A/FUNC.UI&D/010/2021-2024).

## Data Availability

The underlying computer code is available in the GitHub repository https://github.com/zsvargaszilay/exploring_foraging_behaviour_with_computer_vision

## Conflicts of Interest

The authors declare no conflict of interest.

## Notes

https://github.com/zsvargaszilay/exploring_foraging_behaviour_with_computer_vision

### Competing Interest Statement

The authors have declared no competing interest.

## References

Akyon, F. C., Altinuc, S. O., & Temizel, A. (2022). Slicing Aided Hyper Inference and Fine-tuning for Small Object Detection. 2022 IEEE International Conference on Image Processing (ICIP), 966–970. 10.1109/ICIP46576.2022.9897990

Barlow, S. E., & O’Neill, M. A. (2020). Technological advances in field studies of pollinator ecology and the future of e-ecology. Current Opinion in Insect Science, 38, 15–25. 10.1016/j.cois.2020.01.008

Besson, M., Alison, J., Bjerge, K., Gorochowski, T. E., Høye, T. T., Jucker, T., Mann, H. M. R., & Clements, C. F. (2022). Towards the fully automated monitoring of ecological communities. Ecology Letters, 25(12), 2753–2775. 10.1111/ele.14123

Biesmeijer, J. C. (2006). Parallel Declines in Pollinators and Insect-Pollinated Plants in Britain and the Netherlands. Science, 313(5785), 351–354. 10/bwk6j6

Bjerge, K., Geissmann, Q., Alison, J., Mann, H. M. R., Høye, T. T., Dyrmann, M., & Karstoft, H. (2023). Hierarchical classification of insects with multitask learning and anomaly detection. Ecological Informatics, 77, 102278. 10.1016/j.ecoinf.2023.102278

Bjerge, K., Karstoft, H., Mann, H. M. R., & Høye, T. T. (2024). A deep learning pipeline for time-lapse camera monitoring of floral environments and insect populations. 10.1101/2024.04.12.589205

Bjerge, K., Mann, H. M. R., & Høye, T. T. (2022). Real-time insect tracking and monitoring with computer vision and deep learning. Remote Sensing in Ecology and Conservation, 8(3), 315–327. 10.1002/rse2.245

Bjerge, K., Nielsen, J. B., Sepstrup, M. V., Helsing-Nielsen, F., & Høye, T. T. (2021). An Automated Light Trap to Monitor Moths (Lepidoptera) Using Computer Vision-Based Tracking and Deep Learning. Sensors, 21(2), 343. 10.3390/s21020343

Bradski, G. (2000). The OpenCV library. Dr. Dobb’s Journal of Software Tools.

Burkle, L. A., & Alarcón, R. (2011). The future of plant–pollinator diversity: Understanding interaction networks across time, space, and global change. American Journal of Botany, 98(3), 528–538. 10.3732/ajb.1000391

Byers, D. L. (2017). Studying plant–pollinator interactions in a changing climate: A review of approaches. Applications in Plant Sciences, 5(6), 1700012. 10.3732/apps.1700012

Carey, C. C., Ward, N. K., Farrell, K. J., Lofton, M. E., Krinos, A. I., McClure, R. P., Subratie, K. C., Figueiredo, R. J., Doubek, J. P., Hanson, P. C., Papadopoulos, P., & Arzberger, P. (2019). Enhancing collaboration between ecologists and computer scientists: Lessons learned and recommendations forward. Ecosphere, 10(5), e02753. 10.1002/ecs2.2753

Darras, K., Batáry, P., Furnas, B. J., Grass, I., Mulyani, Y. A., & Tscharntke, T. (2019). Autonomous sound recording outperforms human observation for sampling birds: A systematic map and user guide. Ecological Applications, 29(6), e01954. 10.1002/eap.1954

Dicks, L. V., Breeze, T. D., Ngo, H. T., Senapathi, D., An, J., Aizen, M. A., Basu, P., Buchori, D., Galetto, L., Garibaldi, L. A., Gemmill-Herren, B., Howlett, B. G., Imperatriz-Fonseca, V. L., Johnson, S. D., Kovács-Hostyánszki, A., Kwon, Y. J., Lattorff, H. M. G., Lungharwo, T., Seymour, C. L., … Potts, S. G. (2021). A global-scale expert assessment of drivers and risks associated with pollinator decline. Nature Ecology & Evolution, 5(10), Article 10. 10.1038/s41559-021-01534-9

Dwyer, B., Nelson, J., & Hansen, T. (2024). Roboflow (1.0) [Computer software]. https://roboflow.com

Evans, L. J., Smith, K. E., & Raine, N. E. (2017). Fast learning in free-foraging bumble bees is negatively correlated with lifetime resource collection. Scientific Reports, 7(1), 496. 10/f92kvk

Feltham, H., Park, K., & Goulson, D. (2014). Field realistic doses of pesticide imidacloprid reduce bumblebee pollen foraging efficiency. Ecotoxicology, 23(3), 317–323. 10/rct

FFmpeg Developers. (2016). Ffmpeg tool (0.2.6) [Computer software]. http://ffmpeg.org/

Gegear, R. J., Otterstatter, M. C., & Thomson, J. D. (2006). Bumble-bee foragers infected by a gut parasite have an impaired ability to utilize floral information. Proceedings of the Royal Society B: Biological Sciences, 273(1590), 1073–1078. 10/bjwfg4

Geslin, B., Baude, M., Mallard, F., & Dajoz, I. (2014). Effect of local spatial plant distribution and conspecific density on bumble bee foraging behaviour. Ecological Entomology, 39(3), 334–342. 10.1111/een.12106

Ghisbain, G., Thiery, W., Massonnet, F., Erazo, D., Rasmont, P., Michez, D., & Dellicour, S. (2023). Projected decline in European bumblebee populations in the twenty-first century. Nature, 1–5. 10.1038/s41586-023-06471-0

Gill, R. J., Ramos-Rodriguez, O., & Raine, N. E. (2012). Combined pesticide exposure severely affects individual- and colony-level traits in bees. Nature, 491(7422), 105–108. 10/f4cg7z

Gillespie, S. D., & Adler, L. S. (2013). Indirect effects on mutualisms: Parasitism of bumble bees and pollination service to plants. Ecology, 94(2), 454–464. 10.1890/12-0406.1

Godfray, H. C. J., Blacquière, T., Field, L. M., Hails, R. S., Petrokofsky, G., Potts, S. G., Raine, N. E., Vanbergen, A. J., & McLean, A. R. (2014). A restatement of the natural science evidence base concerning neonicotinoid insecticides and insect pollinators. Proceedings of the Royal Society B: Biological Sciences, 281(1786), 20140558. 10.1098/rspb.2014.0558

Goulson, D. (2000). Why do pollinators visit proportionally fewer flowers in large patches? Oikos, 91(3), 485–492. 10.1034/j.1600-0706.2000.910309.x

Goulson, D. (2010). Bumblebees: Behaviour, ecology, and conservation (2nd ed). Oxford University Press.

Goulson, D., Lye, G. C., & Darvill, B. (2008). Decline and Conservation of Bumble Bees. Annual Review of Entomology, 53(1), 191–208. 10.1146/annurev.ento.53.103106.093454

Harder, L. D. (1983). Flower handling efficiency of bumble bees: Morphological aspects of probing time. Oecologia, 57(1), 274–280. 10.1007/BF00379591

Harder, L. D. (1986). Effects of nectar concentration and flower depth on flower handling efficiency of bumble bees. Oecologia, 69(2), 309–315. 10.1007/BF00377639

Harris, C. R., Millman, K. J., Van Der Walt, S. J., Gommers, R., Virtanen, P., Cournapeau, D., Wieser, E., Taylor, J., Berg, S., Smith, N. J., Kern, R., Picus, M., Hoyer, S., Van Kerkwijk, M. H., Brett, M., Haldane, A., Del Río, J. F., Wiebe, M., Peterson, P., … Oliphant, T. E. (2020). Array programming with NumPy. Nature, 585(7825), 357–362. 10.1038/s41586-020-2649-2

Harris, C., & Ratnieks, F. L. W. (2024). Improving pollen and nectar supply by identifying the red clover (*Trifolium pratense*) cultivars that attract most pollinators. Agricultural and Forest Entomology, 26(2), 273–283. 10.1111/afe.12613

Høye, T. T., Ärje, J., Bjerge, K., Hansen, O. L. P., Iosifidis, A., Leese, F., Mann, H. M. R., Meissner, K., Melvad, C., & Raitoharju, J. (2021). Deep learning and computer vision will transform entomology. Proceedings of the National Academy of Sciences, 118(2). 10.1073/pnas.2002545117

Hunter, J. D. (2007). Matplotlib: A 2D graphics environment. Computing in Science & Engineering, 9(3), 90–95. 10.1109/MCSE.2007.55

Jha, S., & Kremen, C. (2013). Resource diversity and landscape-level homogeneity drive native bee foraging. Proceedings of the National Academy of Sciences, 110(2), 555–558. 10.1073/pnas.1208682110

Jocher, G. (2020). *ultralytics/yolov5: V3.1* (v3.1) [Computer software]. Zenodo. 10.5281/zenodo.4154370

Jocher, G., Chaurasia, A., & Qiu, J. (2023). Ultralytics yolov8 (8.0.0) [Computer software]. https://github.com/ultralytics/ultralytics

Kaiser-Bunbury, C. N., Muff, S., Memmott, J., Müller, C. B., & Caflisch, A. (2010). The robustness of pollination networks to the loss of species and interactions: A quantitative approach incorporating pollinator behaviour. Ecology Letters, 13(4), 442–452. 10.1111/j.1461-0248.2009.01437.x

Kerr, J. T., Pindar, A., Galpern, P., Packer, L., Potts, S. G., Roberts, S. M., Rasmont, P., Schweiger, O., Colla, S. R., Richardson, L. L., Wagner, D. L., Gall, L. F., Sikes, D. S., & Pantoja, A. (2015). Climate change impacts on bumblebees converge across continents. Science, 349(6244), 177–180. 10/f7jjqp

Kleijn, D., Winfree, R., Bartomeus, I., Carvalheiro, L. G., Henry, M., Isaacs, R., Klein, A.-M., Kremen, C., M’Gonigle, L. K., Rader, R., Ricketts, T. H., Williams, N. M., Lee Adamson, N., Ascher, J. S., Báldi, A., Batáry, P., Benjamin, F., Biesmeijer, J. C., Blitzer, E. J., … Potts, S. G. (2015). Delivery of crop pollination services is an insufficient argument for wild pollinator conservation. Nature Communications, 6(1), Article 1. 10.1038/ncomms8414

Krell, R. (2018). The pollination of cultivated plants: A compendium for practitioners (Vol. 2). Food and Agriculture Organization of the United Nations.

Lázaro, A., Aase, A. L. T. O., & Totland, Ø. (2011). Relationships between densities of previous and simultaneous foragers and the foraging behaviour of three bumblebee species. Ecological Entomology, 36(2), 221–230. 10.1111/j.1365-2311.2011.01263.x

Lippert, C., Feuerbacher, A., & Narjes, M. (2021). Revisiting the economic valuation of agricultural losses due to large-scale changes in pollinator populations. Ecological Economics, 180, 106860. 10.1016/j.ecolecon.2020.106860

Lövei, G. L., & Ferrante, M. (2024). The Use and Prospects of Nonlethal Methods in Entomology. Annual Review of Entomology, 69(1), 183–198. 10.1146/annurev-ento-120220-024402

Martinet, B., Dellicour, S., Ghisbain, G., Przybyla, K., Zambra, E., Lecocq, T., Boustani, M., Baghirov, R., Michez, D., & Rasmont, P. (2020). Global effects of extreme temperatures on wild bumblebees. Conservation Biology, n/a(n/a). 10/ghqvcg

Nath, R., Singh, H., & Mukherjee, S. (2023). Insect pollinators decline: An emerging concern of Anthropocene epoch. Journal of Apicultural Research, 62(1), 23–38. 10.1080/00218839.2022.2088931

Ngo, T. N., Rustia, D. J. A., Yang, E.-C., & Lin, T.-T. (2021). Automated monitoring and analyses of honey bee pollen foraging behavior using a deep learning-based imaging system. Computers and Electronics in Agriculture, 187, 106239. 10.1016/j.compag.2021.106239

Nieto, A. (2014). European Red List of Bees. Publication Office of the European Union. https://portals.iucn.org/library/node/45219

Ollerton, J., Winfree, R., & Tarrant, S. (2011). How many flowering plants are pollinated by animals? Oikos, 120(3), 321–326. 10.1111/j.1600-0706.2010.18644.x

Paszke, A., Gross, S., Massa, F., Lerer, A., Bradbury, J., Chanan, G., Killeen, T., Lin, Z., Gimelshein, N., Antiga, L., Desmaison, A., Kopf, A., Yang, E., DeVito, Z., Raison, M., Tejani, A., Chilamkurthy, S., Steiner, B., Fang, L., … Chintala, S. (2019). PyTorch: An imperative style, high-performance deep learning library. In Advances in neural information processing systems 32 (pp. 8024–8035). Curran Associates, Inc. http://papers.neurips.cc/paper/9015-pytorch-an-imperative-style-high-performance-deep-learning-library.pdf

Potts, S. G., Biesmeijer, J. C., Kremen, C., Neumann, P., Schweiger, O., & Kunin, W. E. (2010). Global pollinator declines: Trends, impacts and drivers. Trends in Ecology & Evolution, 25(6), 345–353. 10.1016/j.tree.2010.01.007

Preti, M., Verheggen, F., & Angeli, S. (2021). Insect pest monitoring with camera-equipped traps: Strengths and limitations. Journal of Pest Science, 94(2), 203–217. 10.1007/s10340-020-01309-4

Pyke, G. (1984). Optimal Foraging Theory: A Critical Review. *Annual Review of Ecology*, Evolution and Systematic, 15, 523–575. 10.1146/annurev.ecolsys.15.1.523

Python Software Foundation. (2019). Python Language Reference (3.8) [Computer software]. http://www.python.org

R Core Team. (2021). R: A language and environment for statistical computing [Computer software]. https://www.R-project.org/

Rao, S., & Stephen, W. P. (2009). Bumble Bee Pollinators in Red Clover Seed Production. Crop Science, 49(6), 2207–2214. 10.2135/cropsci2009.01.0003

Rasband, W. S. (2018). ImageJ 1997-2018. [Computer software]. U. S. National Institutes of Health. https://imagej.nih.gov/ij/

Ratnayake, M. N., Amarathunga, D. C., Zaman, A., Dyer, A. G., & Dorin, A. (2022). Spatial Monitoring and Insect Behavioural Analysis Using Computer Vision for Precision Pollination (arXiv:2205.04675). arXiv. http://arxiv.org/abs/2205.04675

Ratnayake, M. N., Dyer, A. G., & Dorin, A. (2021). Tracking individual honeybees among wildflower clusters with computer vision-facilitated pollinator monitoring. PLOS ONE, 16(2), e0239504. 10.1371/journal.pone.0239504

Samuelson, E. E. W., Chen-Wishart, Z. P., Gill, R. J., & Leadbeater, E. (2016). Effect of acute pesticide exposure on bee spatial working memory using an analogue of the radial-arm maze. Scientific Reports, 6, 38957. 10/f9f2g2

Schweiger, O., Biesmeijer, J. C., Bommarco, R., Hickler, T., Hulme, P. E., Klotz, S., Kühn, I., Moora, M., Nielsen, A., Ohlemüller, R., Petanidou, T., Potts, S. G., Pyšek, P., Stout, J. C., Sykes, M. T., Tscheulin, T., Vilà, M., Walther, G.-R., Westphal, C., … Settele, J. (2010). Multiple stressors on biotic interactions: How climate change and alien species interact to affect pollination. Biological Reviews, 85(4), 777–795. 10.1111/j.1469-185X.2010.00125.x

Sittinger, M., Uhler, J., Pink, M., & Herz, A. (2024). Insect detect: An open-source DIY camera trap for automated insect monitoring. PLOS ONE, 19(4), e0295474. 10.1371/journal.pone.0295474

Somme, L., Vanderplanck, M., Michez, D., Lombaerde, I., Moerman, R., Wathelet, B., Wattiez, R., Lognay, G., & Jacquemart, A.-L. (2015). Pollen and nectar quality drive the major and minor floral choices of bumble bees. Apidologie, 46(1), 92–106. 10.1007/s13592-014-0307-0

Soroye, P., Newbold, T., & Kerr, J. (2020). Climate change contributes to widespread declines among bumble bees across continents. Science, 367(6478), 685–688. 10/dk78

Spiesman, B. J., Gratton, C., Hatfield, R. G., Hsu, W. H., Jepsen, S., McCornack, B., Patel, K., & Wang, G. (2021). Assessing the potential for deep learning and computer vision to identify bumble bee species from images. Scientific Reports, 11(1), 7580. 10.1038/s41598-021-87210-1

Stanley, D. A., Garratt, M. P. D., Wickens, J. B., Wickens, V. J., Potts, S. G., & Raine, N. E. (2015). Neonicotinoid pesticide exposure impairs crop pollination services provided by bumblebees. Nature, 528(7583), 548–550. 10/f7373p

Stanley, D. A., Russell, A. L., Morrison, S. J., Rogers, C., & Raine, N. E. (2016). Investigating the impacts of field-realistic exposure to a neonicotinoid pesticide on bumblebee foraging, homing ability and colony growth. Journal of Applied Ecology, 53(5), 1440–1449. 10/f9csvr

Steen, R. (2017). Diel activity, frequency and visit duration of pollinators in focal plants: In situ automatic camera monitoring and data processing. Methods in Ecology and Evolution, 8(2), 203–213. 10.1111/2041-210X.12654

Stout, J. C., Allen, J. A., & Goulson, D. (1998). The influence of relative plant density and floral morphological complexity on the behaviour of bumblebees. Oecologia, 117(4), 543–550. 10.1007/s004420050691

Stout, J. C., & Goulson, D. (2002). The influence of nectar secretion rates on the responses of bumblebees (Bombus spp.) to previously visited flowers. Behavioral Ecology and Sociobiology, 52(3), 239–246. 10.1007/s00265-002-0510-2

Szabo, T. I., & Najda, H. G. (1985). Flowering, Nectar Secretion and Pollen Production of Some Legumes in the Peace River Region of Alberta, Canada. Journal of Apicultural Research, 24(2), 102–106. 10.1080/00218839.1985.11100656

The pandas development team. (2022). pandas-dev/pandas: Pandas (v1.5.2) [Computer software]. Zenodo. 10.5281/zenodo.7344967

Treanore, E., Barie, K., Derstine, N., Gadebusch, K., Orlova, M., Porter, M., Purnell, F., & Amsalem, E. (2021). Optimizing Laboratory Rearing of a Key Pollinator, Bombus impatiens. Insects, 12(8), Article 8. 10.3390/insects12080673

Tuia, D., Kellenberger, B., Beery, S., Costelloe, B. R., Zuffi, S., Risse, B., Mathis, A., Mathis, M. W., Van Langevelde, F., Burghardt, T., Kays, R., Klinck, H., Wikelski, M., Couzin, I. D., Van Horn, G., Crofoot, M. C., Stewart, C. V., & Berger-Wolf, T. (2022). Perspectives in machine learning for wildlife conservation. Nature Communications, 13(1), 792. 10.1038/s41467-022-27980-y

Vaca-Uribe, J. L., Figueroa, L. L., Santamaría, M., & Poveda, K. (2021). Plant richness and blooming cover affect abundance of flower visitors and network structure in Colombian orchards. Agricultural and Forest Entomology, 23(4), 545–556. 10.1111/afe.12460

van Klink, R., August, T., Bas, Y., Bodesheim, P., Bonn, A., Fossøy, F., Høye, T. T., Jongejans, E., Menz, M. H. M., Miraldo, A., Roslin, T., Roy, H. E., Ruczyński, I., Schigel, D., Schäffler, L., Sheard, J. K., Svenningsen, C., Tschan, G. F., Wäldchen, J., … Bowler, D. E. (2022). Emerging technologies revolutionise insect ecology and monitoring. Trends in Ecology & Evolution, 37(10), 872–885. 10.1016/j.tree.2022.06.001

Vanbergen, A. J. (2014). Landscape alteration and habitat modification: Impacts on plant-pollinator systems. Current Opinion in Insect Science, 5, 44–49. 10.1016/j.cois.2014.09.004

Vanbergen, A. J., & Initiative, the I. P. (2013). Threats to an ecosystem service: Pressures on pollinators. Frontiers in Ecology and the Environment, 11(5), 251–259. 10.1890/120126

Varga-Szilay, Z. (2023a). Persicaria_capitata_2022 Dataset [Open Source Dataset]. Roboflow. https://universe.roboflow.com/zsofia-varga-szilay/persicaria_capitata_20220921_22

Varga-Szilay, Z. (2023b). Lotus_creticus_2022 Dataset [Open Source Dataset]. Roboflow. https://universe.roboflow.com/zsofia-varga-szilay/lotus_creticus_2022

Varga-Szilay, Z. (2023c). Trifolium_pratense_2022 Dataset [Open Source Dataset]. Roboflow. https://universe.roboflow.com/zsofia-varga-szilay/trifolium_pratense_2022

Varga-Szilay, Z., & Tóth, Z. (2022). Is acetamiprid really not that harmful to bumblebees (Apidae: Bombus spp.)? Apidologie, 53(1), 2. 10.1007/s13592-022-00909-6

Velthuis, H. H. W., & van Doorn, A. (2006). A century of advances in bumblebee domestication and the economic and environmental aspects of its commercialization for pollination. Apidologie, 37(4), 421–451. 10.1051/apido:2006019

Virtanen, P., Gommers, R., Oliphant, T. E., Haberland, M., Reddy, T., Cournapeau, D., Burovski, E., Peterson, P., Weckesser, W., Bright, J., van der Walt, S. J., Brett, M., Wilson, J., Millman, K. J., Mayorov, N., Nelson, A. R. J., Jones, E., Kern, R., Larson, E., … SciPy 1.0 Contributors. (2020). SciPy 1.0: Fundamental algorithms for scientific computing in python. Nature Methods, 17, 261–272. 10.1038/s41592-019-0686-2

Wickham, H. (2016). ggplot2: Elegant graphics for data analysis. Springer-Verlag New York. https://ggplot2.tidyverse.org

Wickham, H., François, R., Henry, L., Müller, K., Vaughan, D., Software, P., & PBC. (2023). dplyr: A Grammar of Data Manipulation (1.1.4) [Computer software]. https://cran.r-project.org/web/packages/dplyr/index.html

Wickham, H., & Henry, L. (2023). purrr: Functional programming tools [Manual]. https://purrr.tidyverse.org/

Wickham, H., Hester, J., & Bryan, J. (2024). readr: Read rectangular text data [Manual]. https://readr.tidyverse.org

Williams, P. H., & Osborne, J. L. (2009). Bumblebee vulnerability and conservation world-wide. Apidologie, 40(3), 367–387. 10/d8s24c

Zeileis, A., & Hothorn, T. (2002). Diagnostic checking in regression relationships. R News, 2(3), 7–10.

Zeileis, A., Köll, S., & Graham, N. (2020). Various versatile variances: An object-oriented implementation of clustered covariances in R. Journal of Statistical Software, 95(1), 1–36. 10.18637/jss.v095.i01

Zimmerman, M. (1981). Optimal foraging, plant density and the marginal value theorem. Oecologia, 49(2), 148–153. 10.1007/BF00349181

