## Supplementary Materials for "Flower visitation through the lens: Exploring the foraging behaviour of *Bombus terrestris* with a computer vision-based application"

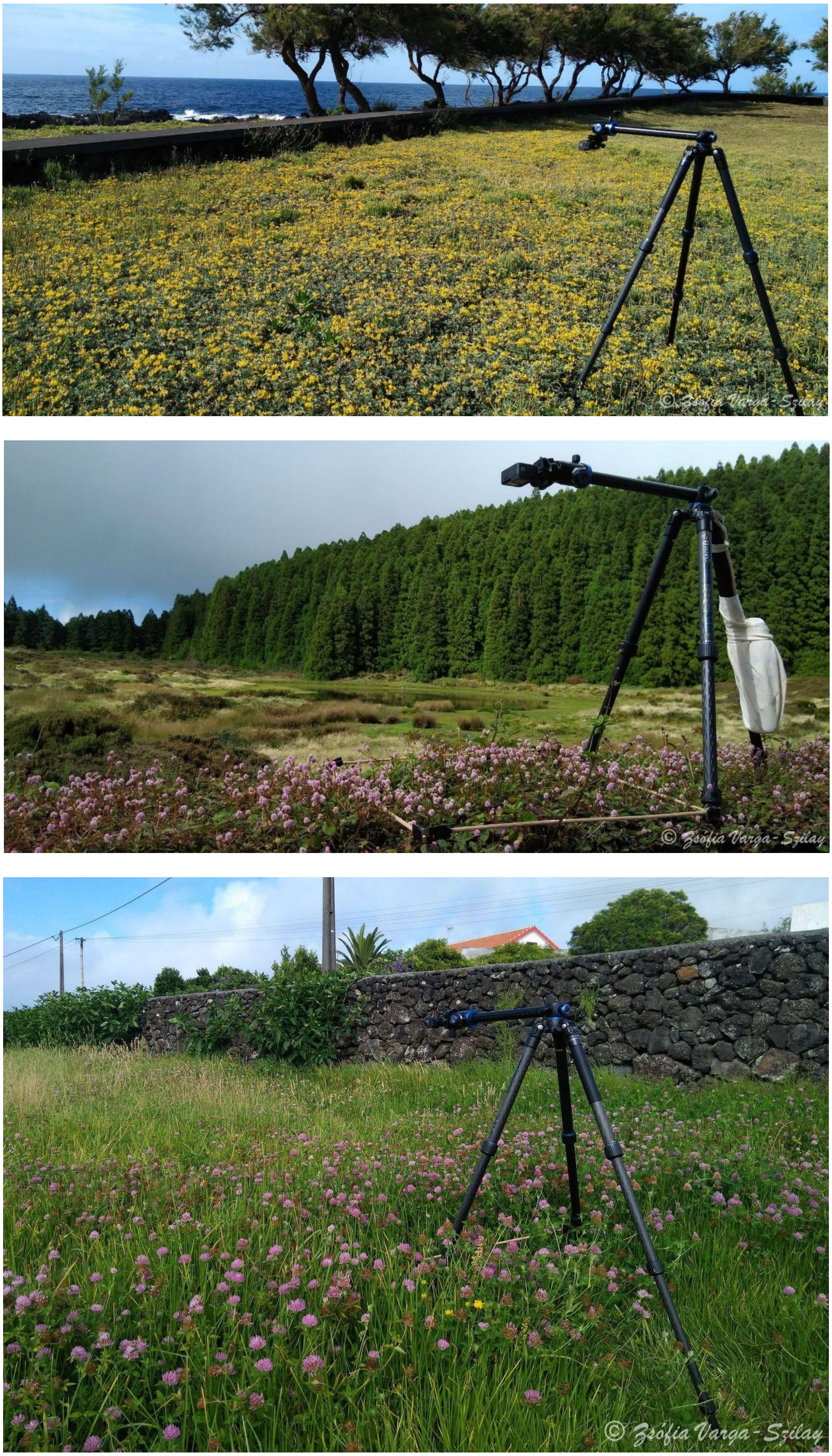


(a) (b) (c)

**Figure S1**: The sampling sites were located in urban areas in Terceira (Azores, Portugal) (a) *Lotus*: 38°48'06.0" N, 27°15'17.8" W at an elevation of 10m a.s.l., (b) *Persicaria*: 38°44'14.7" N, 27°16'07.8" W 548m a.s.l., and (c) *Trifolium*: 38°47'38.7" N, 27°15'24.0" W 52m a.s.l..

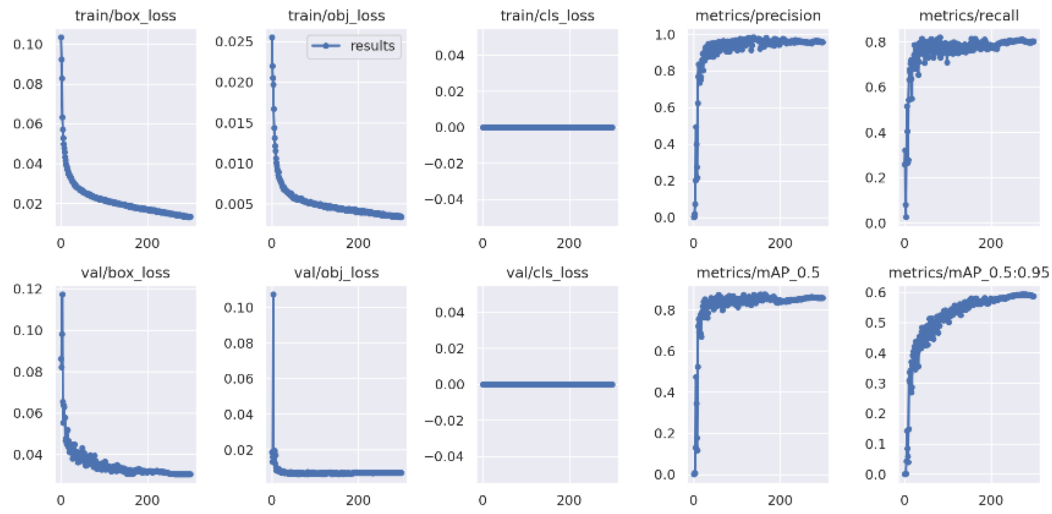


(a)

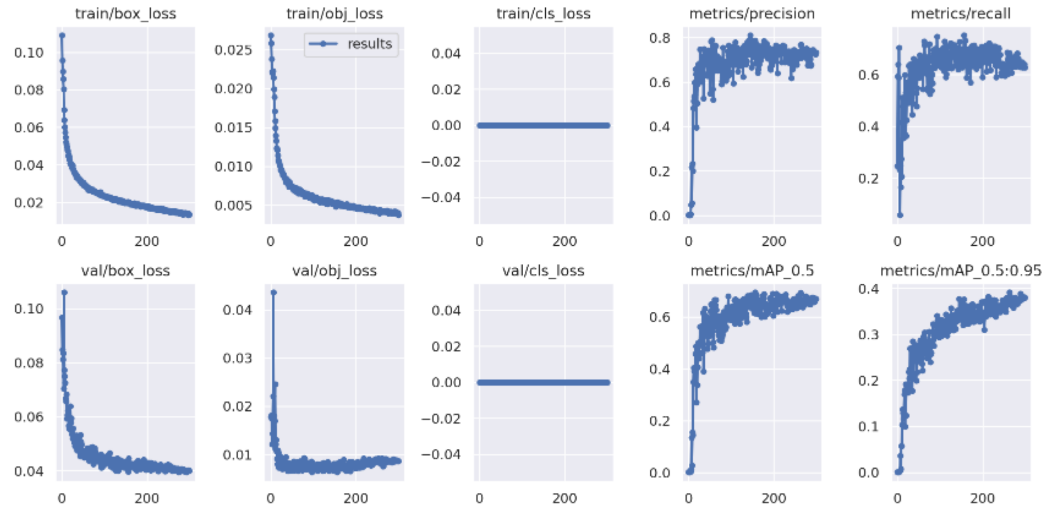


(b)

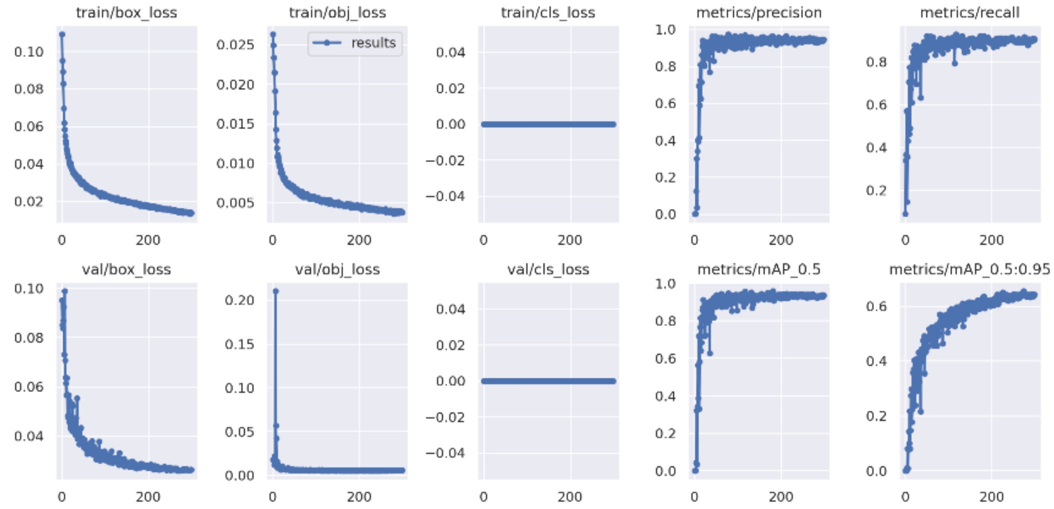


(c)

**Figure S2**: Specific evaluation metrics of YOLOv5 models (trained without pre-trained weights) on our three datasets: (a) *Lotus creticus*, (b) *Persicaria capitata*, and (c) *Trifolium pratense*. The change in key indicators, including *box_loss* (bounding box regression loss), *obj_loss* (confidence of object presence), *cls_loss* (classification loss, with 'bumblebee' as the only class), *precision, recall, and mAP* (mean Average Precision), over the course of training (300 epochs).

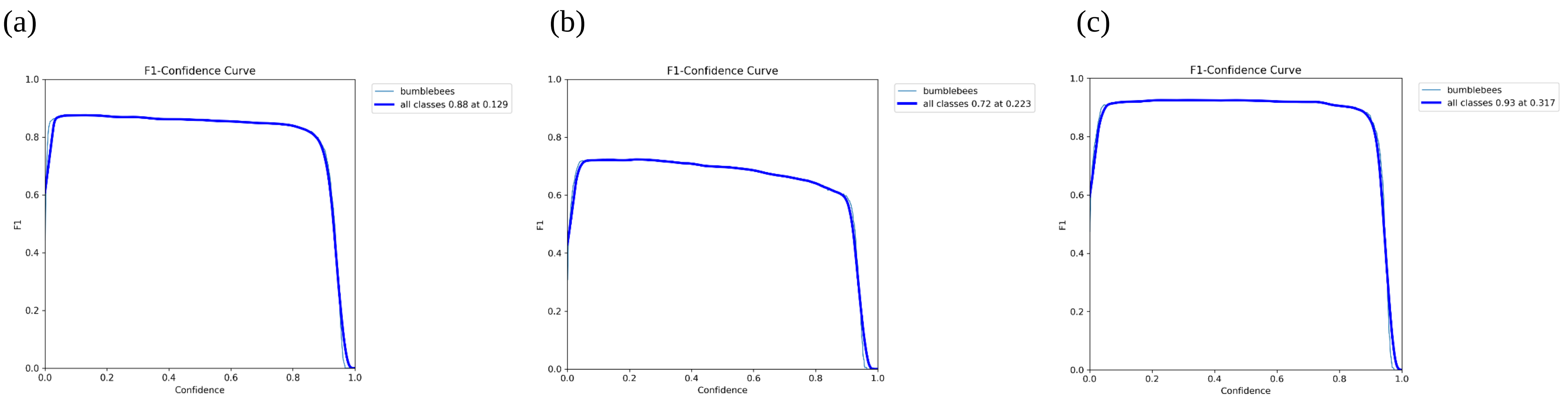


**Figure S3:** F1 scores of the trained YOLOv5 model demonstrating the balance between precision and recall in detecting bumblebees on (a) *Lotus creticus*, (b) *Persicaria capitata*, and (c) *Trifolium pratense*.

**Table S1**: F1 scores after post-processing, based on the model predictions for 30 randomly selected frames from 5 random videos per plant species. The formula used for the F1 score calculation: F1 = 2 TP / (2 TP + FP + FN).

| | | Predicted | |
|---|---|---|---|
| | | **Positive** (Bumblebee) | **Negative** (Bumblebee) |
| Observed | **Positive** (Bumblebee) | **True positive (TP)** *Lotus*: 25 *Persicaria*: 21 *Trifolium*: 20 | **False negative (FN)** *Lotus*: 2 *Persicaria*: 0 *Trifolium: 2* |
| | **Negative** (No bumblebee) | **False positive (FP)** *Lotus*: 5 *Persicaria*: 6 *Trifolium*: 0 | |
| | | | **F1 score** *Lotus*: **0.88** *Persicaria*: **0.88** *Trifolium*: **0.95** |

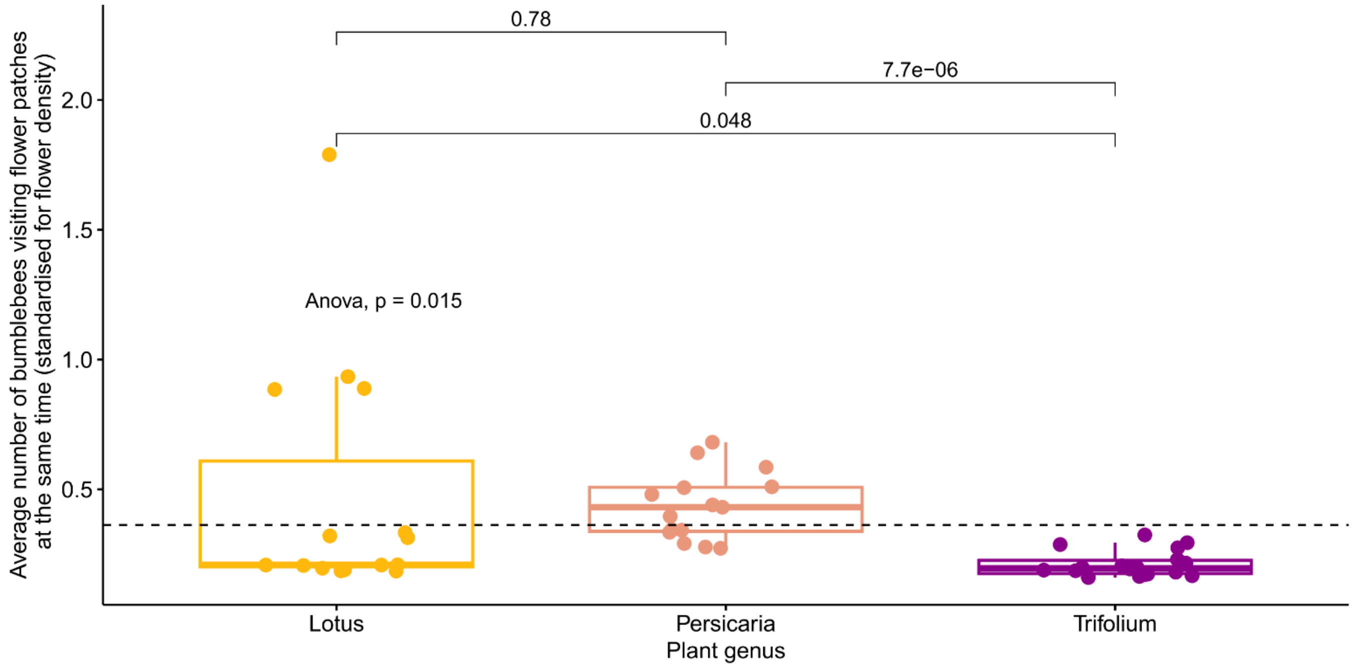


**Figure S4:** The average number of bumblebees visiting flower patches at the same time (standardised for flower cover) separated by plant species. The global p-value for the ANOVA test is shown in the figure, as are the pairwise comparisons (t-tests) of the averages between plant species. The dashed line shows the mean of the y-axis. Each point represents one video (n = 15, 15, and 18, for *Lotus*, *Persicaria*, and *Trifolium*, respectively).
